# Lineage-specific neutralising antibodies after SARS-CoV-2 mild disease. Immune boosting effect of vaccination

**DOI:** 10.1101/2024.11.08.622599

**Authors:** Carlos Davina-Nunez, Sonia Perez-Castro, Jorge-Julio Cabrera-Alvargonzalez, Elena Gonzalez-Alonso, Sergio Silva-Bea, Miriam Rodriguez-Perez, Maria del Pilar Figueroa-Lamas, Alexandre Perez-Gonzalez, Víctor del Campo, Almudena Rojas, Joaquin Mendoza, Benito Regueiro-Garcia

## Abstract

We followed a group of 105 non-vaccinated individuals after Alpha or Delta SARS-CoV-2 mild disease, measuring the viral shedding (qRT-PCR, dPCR and subgenomic RNA-E) and humoral response (commercial immunoassay and pseudovirus and live virus neutralisation) up to six months. Sixty nine patients received a vaccination boost during the follow-up period (n=95). Subgenomic RNA-E showed a shorter period until negativity (mean 2.2 weeks) compared to gRNA (mean 5.2 weeks). A high correlation between qRT-PCR and dPCR was found for viral load estimation, even when no nucleic acid extraction was used in dPCR (R^2^ = 0.87). Post-convalescent sera showed the strongest neutralisation against the variant of natural exposure, while the neutralisation capacity against Omicron variants was significantly lower compared to the other variants. Additionally, the results suggested that commercial immunoassays may not accurately predict protection against a different variant than the variant of exposure. An immune boosting effect of the SARS-CoV-2 vaccination was evident.

Variant-specific neutralising antibodies were detected one month after natural infection. Although short lived, maximum igG response was observed after hybrid immunisation (natural infection + vaccination). This study also points to potential improvements in the clinical management of SARS-CoV-2 cases. Firstly, subgenomic RNA-E is a potentially more accurate biomarker of infectivity than current qRT-PCRs using genomic RNA as target. Secondly, accuracy of high-throughput immunoassays must be validated in order to estimate specific protection and organise vaccination campaigns. Our findings could play a role in the current implementation of SARS-CoV-2 vaccine programs.

**Author Summary:** Years after SARS-CoV-2 related infections challenged the healthcare systems of the whole world, the optimal strategy to deal with diagnosis, quarantines or vaccination patterns is still a matter of debate. The interplay between infectivity and immunity in the different circulating variants is complex, and qPCR for diagnosis may extend quarantines, as detection of viral RNA does not necessarily mean that the virus remains infectious.

We studied a group of patients infected with pre-Omicron variants to study how their variant-specific antibodies reacted to the past and present variants of SARS-CoV-2. In both Alpha and Delta, antibodies neutralise their own variants better than other variants that came before and after. Perhaps most importantly, neutralisation was lowest against Omicron variants.

We followed the viral shedding dynamics of the patients, testing different PCR techniques and targets. Subgenomic RNA was detectable in nasopharyngeal samples for a shorter time than genomic RNA and it has been suggested as a good marker of infectivity. Using sgRNA instead of genomic RNA as a PCR target could reduce hospital bed occupation and quarantine time. Overall, we hope that these results could help guide pandemic and diagnostic control in the future.

## Introduction

Having an accurate knowledge on SARS-CoV-2 viral shedding dynamics and neutralising immunity is key in the control of viral transmission and vaccination campaign organisation. However, despite all the findings about SARS-CoV-2 dynamics since the discovery of this pathogen, there are still insights to be obtained. For example, most studies have been focused only on hospitalised patients [1]. Viral shedding in mild COVID has been therefore understudied in comparison, despite being the majority of the infected individuals in a community.

The RT-PCR from a nasopharyngeal swab remains the gold standard for SARS-CoV-2 diagnosis. It must be noted that, in this clinical specimen, viral load in a sample can be more or less representative of the viral load in the patient, as variations in the swabbing technique could affect the result [2]. Additionally, the main limitation of this technique is its inability to distinguish between infectious and non-infectious RNA [3]. This is relevant, as it has been previously established that the period of detectable viral RNA is longer than the period of infectivity [1]. Relying only in RT-PCR data could lead to unnecessary extended quarantines, putting excessive pressure in healthcare systems and in individuals undergoing self-isolation.

Viral culture allows distinguishing infectious from non-infectious samples, but it is unrealistic to establish culture as a routine diagnostic technique as it is more expensive and requires a BSL-3 laboratory to grow the virus. As a second limitation, when facing a new virus optimal culture conditions are not known, which could limit the accuracy of viral culture as a measurement of viral infectivity at the early stages of facing a new pathogen [4]. Finding a good biomarker for viral infectivity is increasingly challenging as the infectious period may be affected by disease severity [5,6], variant [3], age [7], gender [8] or assay used [9].

Alternative biomarkers of active infection have been tested, such as the detection of subgenomic RNAs (sgRNAs). Since these molecules are necessary for viral protein translation and generated only during viral replication, they should be only present during periods of active infection [10]. However, more data is required to understand the potential role of sgRNA RT-PCR as a marker of active infection.

An alternative tool for viral RNA quantification is the digital PCR (dPCR). Since the beginning of the SARS-CoV-2 pandemic, dPCR has been proposed as a method for improved detection in low viral load samples [11]. Additionally, it could potentially work reliably even without an RNA extraction process [12], which would decrease processing sample time and improve reproducibility.

The gold standard for measuring humoral immunity *in vitro* is the neutralising antibody assay, and it has provided prediction of protection since the beginning of the SARS-CoV-2 pandemic [13]. However, these assays are challenging to perform in the routine clinical surveillance setting due to limitations in cost, expertise and the biosafety levels required to operate live viruses [14,15].

As an alternative, *in vitro* serological tests are cheap, high-throughput, and can provide models to help understand the immunity in big population groups [16]. These tests could also reduce the demands of large trials in vaccine development by using antibody values to measure protection levels (immuno-bridging) [17]. However, serological tests require standardisation [18] and cannot differentiate the effectiveness of different antibody fractions against different variants. Therefore, neutralising antibody assays (nAb) can achieve a more accurate measurement of protection, especially against new variants of exposure.

Since the beginning of the SARS-CoV-2 pandemic, vaccination campaigns have been organised against variants wild-type, BA.4-5 and XBB.1.5 [19]. Early vaccines against the Wuhan wild-type variant showed a decreased neutralising antibody response against Delta and Omicron variants [20,21], therefore prompting the development of vaccines adapted to the new variants. Measuring the impact of booster vaccinations is challenging however, as a growing number of the population has hybrid immunity as a consequence of breakthrough infections [22].

This study intends to shed light on the viral dynamics along the pre-Omicron SARS-CoV-2 variants of concern (VOCs) mild disease in unvaccinated individuals, as well as the evolution of their post-infection immune response to understand the role of the specific variant of first exposure (mainly Alpha or Delta) on their humoral response.

## Results

A total of 105 patients were recruited for follow-up in either viral excretion dynamics (group 1, 73 Alpha, 10 Delta, 22 others) or IgG levels after the infection (group 2, 67 Alpha, 9 Delta, 19 others). The description of the population can be found in Table 1.

**Table 1.**
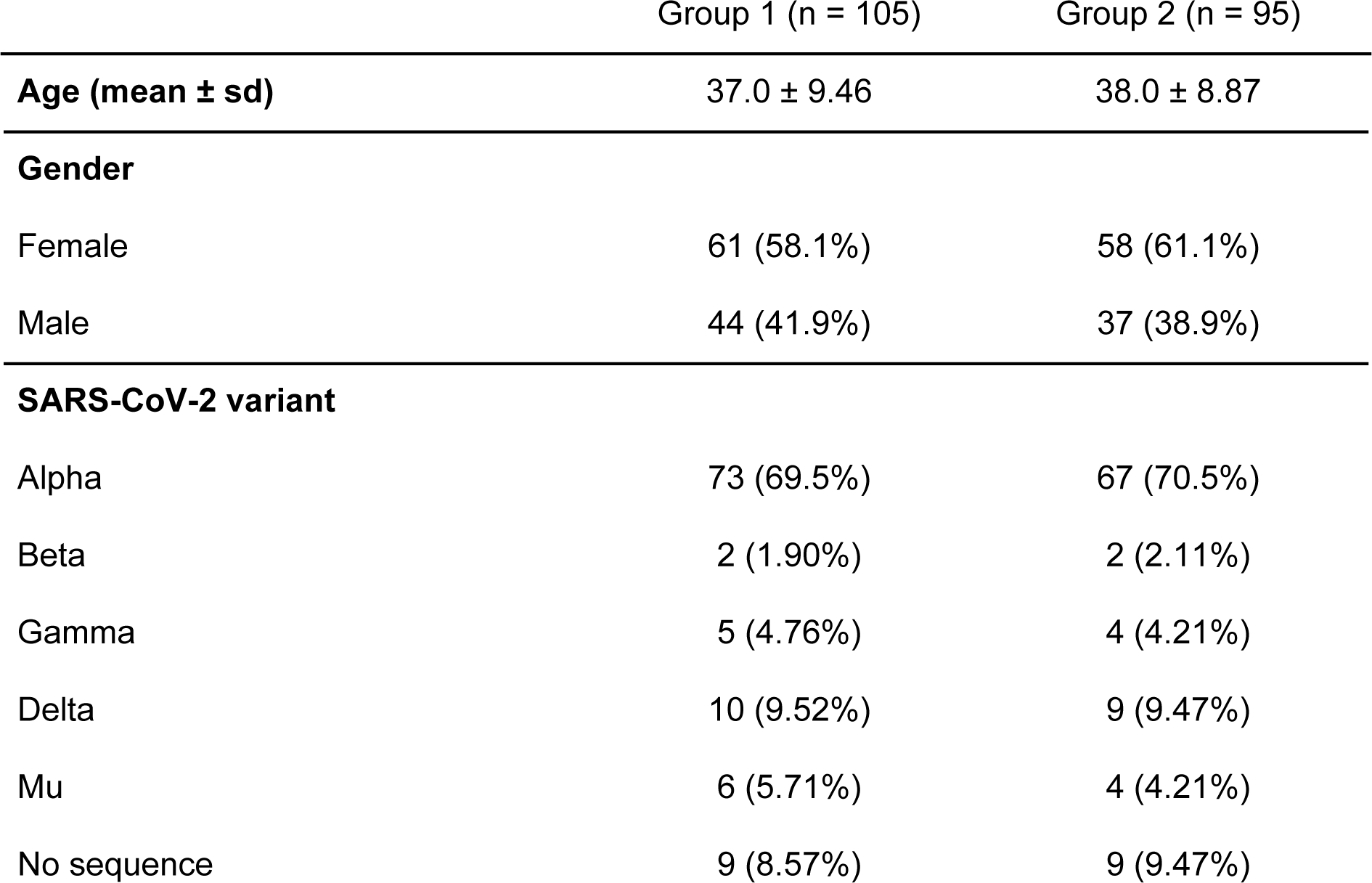
Descriptive of the population participating in the study. Group 1: excretion dynamics. Group 2: humoral immunity.

| Group 1 (n = 105) | Group 2 (n = 95) |
| --- | --- |

|  |  |  |
| --- | --- | --- |
| **Age (mean ± sd)** | 37.0 ± 9.46 | 38.0 ± 8.87 |
| **Gender** |  |  |
| Female | 61 (58.1%) | 58 (61.1%) |
| Male | 44 (41.9%) | 37 (38.9%) |
| **SARS-CoV-2 variant** |  |  |
| Alpha | 73 (69.5%) | 67 (70.5%) |
| Beta | 2 (1.90%) | 2 (2.11%) |
| Gamma | 5 (4.76%) | 4 (4.21%) |
| Delta | 10 (9.52%) | 9 (9.47%) |
| Mu | 6 (5.71%) | 4 (4.21%) |
| No sequence | 9 (8.57%) | 9 (9.47%) |

### Correlation between RT-PCR results and dPCR is solid in results below CT 30

All samples with a positive RT-PCR result from RNA extract in the study were quantified using QiaAcuity digital PCR (Qiagen, Hilgen, Germany) and crude samples as input (direct quantification). Prior to the assays, direct quantification by dPCR was compared to RNA extract to confirm comparable results (S1 Fig), as it has been previously analysed in another study [12].

Results of dPCR showed high correlation with RT-PCR positive samples (R^2^ = 0,84; Fig 1A). This correlation was high for samples below CT 30 (R^2^ = 0.87) (Fig 1B), and negligible in samples with a CT value over 30 (R^2^ = 0.08), showing high uncertainty in viral load determination in samples of low viral load. According to the linear model (Fig 1B), a Ct of 30 is equivalent to 6-7 molecules per microliter in the dPCR data.

**Fig 1.**
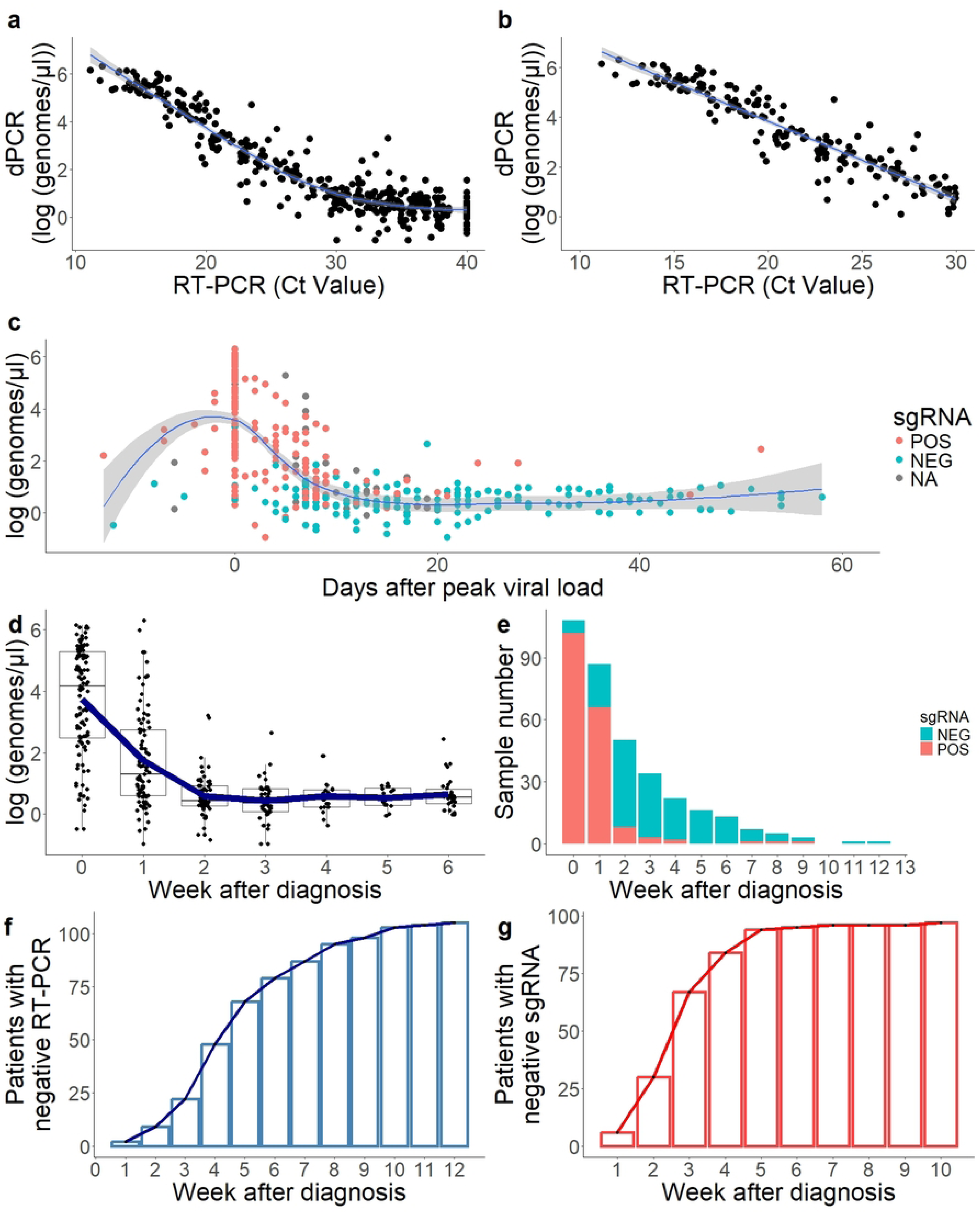
Study of SARS-CoV-2 viral excretion dynamics. **a, b)** Correlation of RT-PCR (ct value) and dPCR (absolute quantification) in estimation of viral load. Correlation is linear until Ct = 30, as shown on the panel on the right. **c)** Excretion dynamics from all positive samples in the study. Day 0 is marked for all subjects as the day of the PCR with the highest viral load. Blue line: LOESS approximation with grey area showing the 95% confidence interval. Point colour indicates the detection of sgRNA in each sample, with NA indicating samples that were not tested for sgRNA. **d)** Boxplot of viral load of all samples per week after diagnosis. The blue line indicates the mean value per week. Six (6) refers to six or more weeks in this plot. **e)** Amount of samples positive in gRNA that were positive in sgRNA (red) or negative (blue) per week after diagnostic. **f)** Amount of patients with a negative RT-PCR in gRNA per week. **g)** Amount of patients with a negative RT-PCR in sgRNA per week. All data used to generate this figure is available in S1 File.

### Most of the convalescent period with detectable viral RNA is a low viral concentration period

In order to get a more realistic grip of infection dynamics, all samples were ordered by the day with detection of maximum viral load. This allowed to normalise all patients according to their infection dynamics, with samples from patients still in the early phases of infection (detection before maximum viral load) and samples from patients in late phases of infection (detection after maximum viral load). There was a period of 20 days with results above 10 genomes/ul, showing a bell curve graph (Fig 1C). After the second week post-infection, a second period was found where RNA is still detectable in most patients but viral load is residual, from an average of 10^3,73^ genomes/μl at time of diagnosis to 10^1,75^ at week 1 and 10^0,56^ at week 2 (Fig 1D). Only seventeen patients (16.2%) showed viral loads below CT 30 from week 2 onwards, and only four (3.8%) from week 3 onwards. However, in 54.3% of the individuals (57/105), viral RNA was still detectable after week 4 after diagnosis (Fig 1F). These late-time positive samples were of low concentration and, considering the data above, challenging to quantify accurately with both qPCR and dPCR.

### sgRNA was only detectable in the first weeks of infection in most samples

Subgenomic RNA, which has been used in previous studies as a marker of viral infectivity [26], was tested in 97 out of 105 patients for sgRNA. Negative sgRNA samples appeared more frequently in the latest days of detectable shedding, with the lowest RNA viral loads (Fig. 1C). When measuring genomic RNA, mean weeks until a negative test (no positive target) was 5.2 ± 2.4 weeks, while with sgRNA it was 2.2 ± 1.4 weeks.

As it is the case with the samples with high viral load, a majority of the samples positive in sgRNA occurred in the weeks 0-1 after diagnosis. Positivity ratio in sgRNA (amongst positive samples as detected by traditional RT-PCR ) was 94.4% (102/108) in the week of diagnosis, 75.8% (66/87) in the following week, and 16.0% (8/50) in the second week, dropping below 10% in the weeks onwards (Fig. 1E). Therefore, starting the second week after diagnosis, only 10.5% of the samples with detectable RNA had detectable sgRNA. Days after diagnosis was a good predictor of sgRNA positivity, with a positivity of 84.9% (169/199) in the first 10 days, and 10.1% (15/148) in the following days until undetectable gRNA. At week 3, 69.1% (67/97) of patients already had a negative result in sgRNA (Fig. 1G).

Notably, one participant tested negative in sgRNA in week 3 and then showed detectable sgRNA back again two months after infection, with its genomic RNA Ct value rising to 24.6 in this period. NGS data confirmed that this participant had the same SARS-CoV-2 strain throughout all the follow-up period, making it unlikely that these were two separate cases of infection.

### Age has a moderate impact in peak viral load, while excretion days remains unchanged

Variables with impact in peak viral load and days of excretion were checked in the population. Age had a mild impact in peak viral load, with older people having a higher viral load (ρ = 0.257, *p* = 0.008; Spearman’s rank correlation coefficient) (S2 FigA,B). The number of excretion days was not significantly affected (ρ = 0.153, *p* = 0.120) (S2 FigA,D). Gender had no impact in excretion dynamics, nor in peak viral load nor in excretion days (*p* = 0.367; *p* = 0.194 respectively; Wilcoxon Signed-rank test) (S2 FigC,E).

### IgG levels reach their peak one month after infection

Sera samples were taken from 95 participants in order to measure variations in the IgG levels after infection in a 6-month period. Given that the follow-up time of this study overlapped with the SARS-CoV-2 vaccination campaign in Spain, it was an opportunity to study the impact of the vaccination after the natural infection. The amount of participants vaccinated at each timepoint is described in table 2.

**Table 2.**
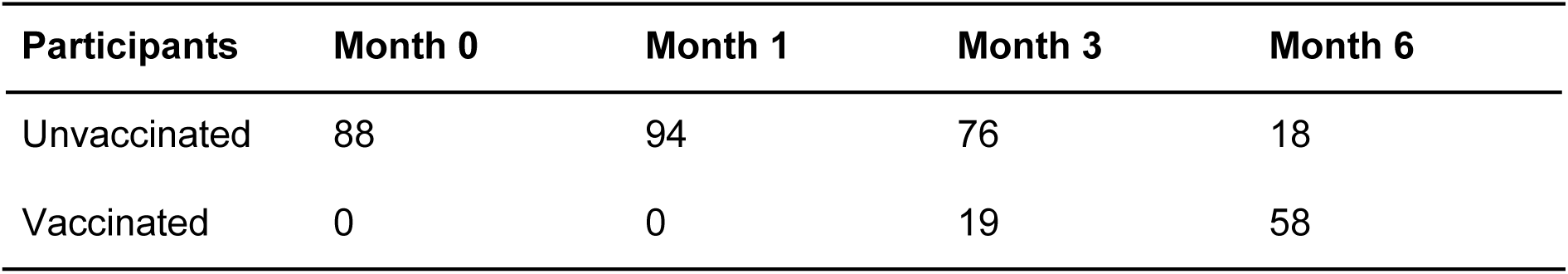
individuals vaccinated across the follow-up vigilance period. All subjects who were vaccinated had only one dose after infection.

| Participants | Month 0 | Month 1 | Month 3 | Month 6 |
| --- | --- | --- | --- | --- |
| Unvaccinated | 88 | 94 | 76 | 18 |
| Vaccinated | 0 | 0 | 19 | 58 |

Unvaccinated individuals throughout the whole follow-up period had a peak of antibody levels at month 1. Although there was a drop of antibody count at months 3 and 6, the difference was non-significant (Fig 2, S2 File, Dunn’s test).

**Fig 2.**
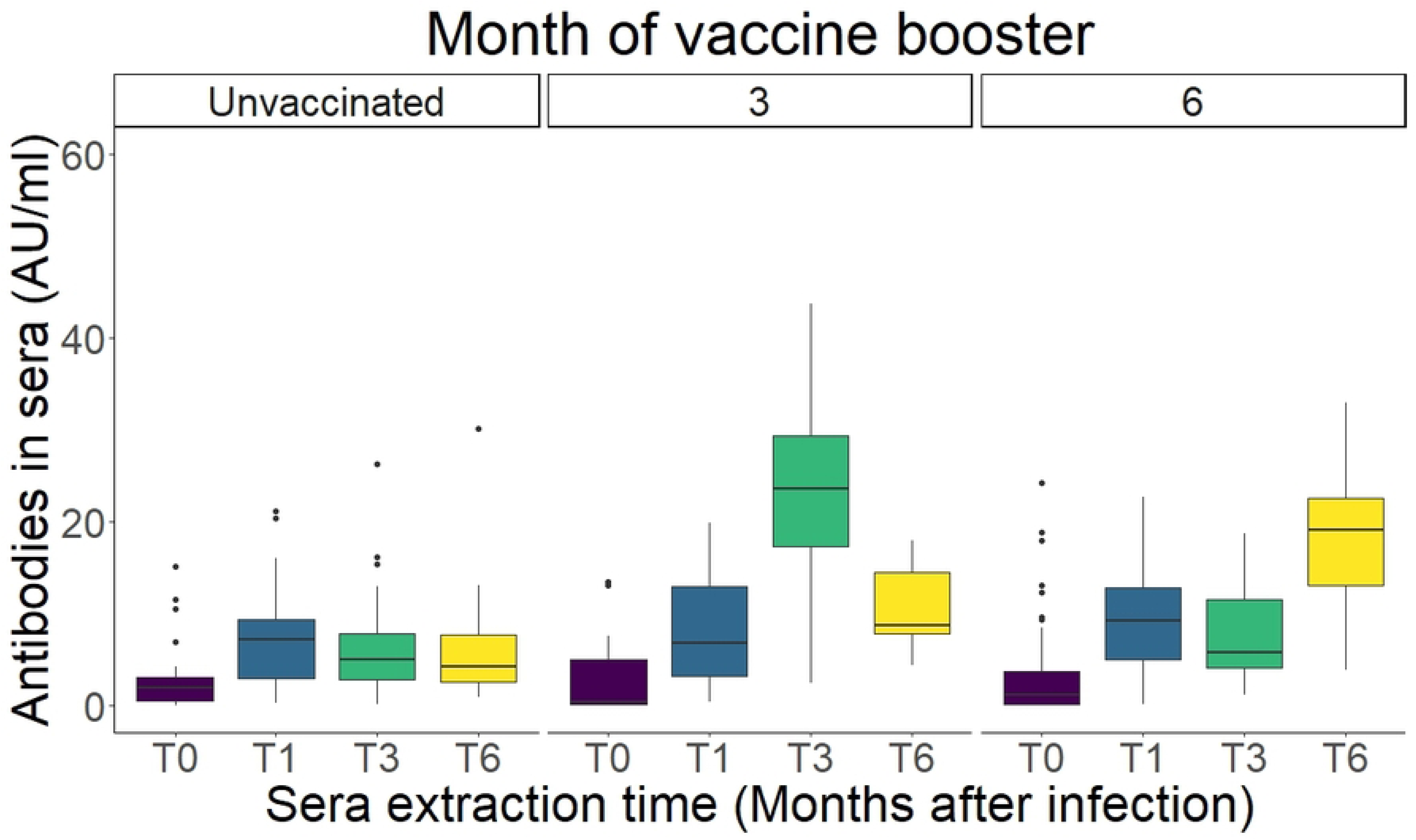
IgG levels of patients according to vaccination time. **Left:** unvaccinated individuals. After a peak in month 1, antibody AUs decrease moderately in months 3 and 6. **Middle:** individuals vaccinated before month 3. A boost of igG from vaccination is shown. However, at month 6 antibody levels already drop to around pre-vaccination levels (month 1). **Right:** individuals vaccinated before month 6. Antibody boost caused by vaccination at the month 6 measurement. All data used to generate this figure is available in S3 File.

Participants showed a boost in antibody levels in the sample immediately after vaccination, with month 3 showing the highest levels measured in the patients with their vaccine taken immediately between months one and three (Fig 2). Nonetheless, at month 6, antibody counts return to no significant difference with the antibody values at month 1 (pre-vaccination) (*p* = 0.187, S2 File). Participants receiving a vaccine dose between months 3 and 6 also showed an increase of antibody levels at month 6, after a non-significant drop between months 1 and 3 (*p* = 0.187, S2 File).

### Stronger neutralisation capacity against the variant of infection

Samples from one month after diagnosis were selected for neutralisation assays, as in this group there were no vaccinated individuals. Individuals were selected from those infected with Alpha and Delta variants, the two most common variants at time of recruitment.

Nine participants were infected with the Delta variant. In order to compare anti-Alpha and anti-Delta sera, 18 samples were selected for the pseudovirus neutralisation assay. In addition to the nine patients with anti-Delta sera fractions, nine participants with anti-Alpha sera were tested. Samples were selected in order to have paired antibody values between Alpha and Delta as measured by an *in vitro* immunoassay (Fig 3A). Neutralisation was performed against a Delta-spike carrying pseudovirus and measured as 50% inhibitory dilution (ID50).

**Fig 3.**
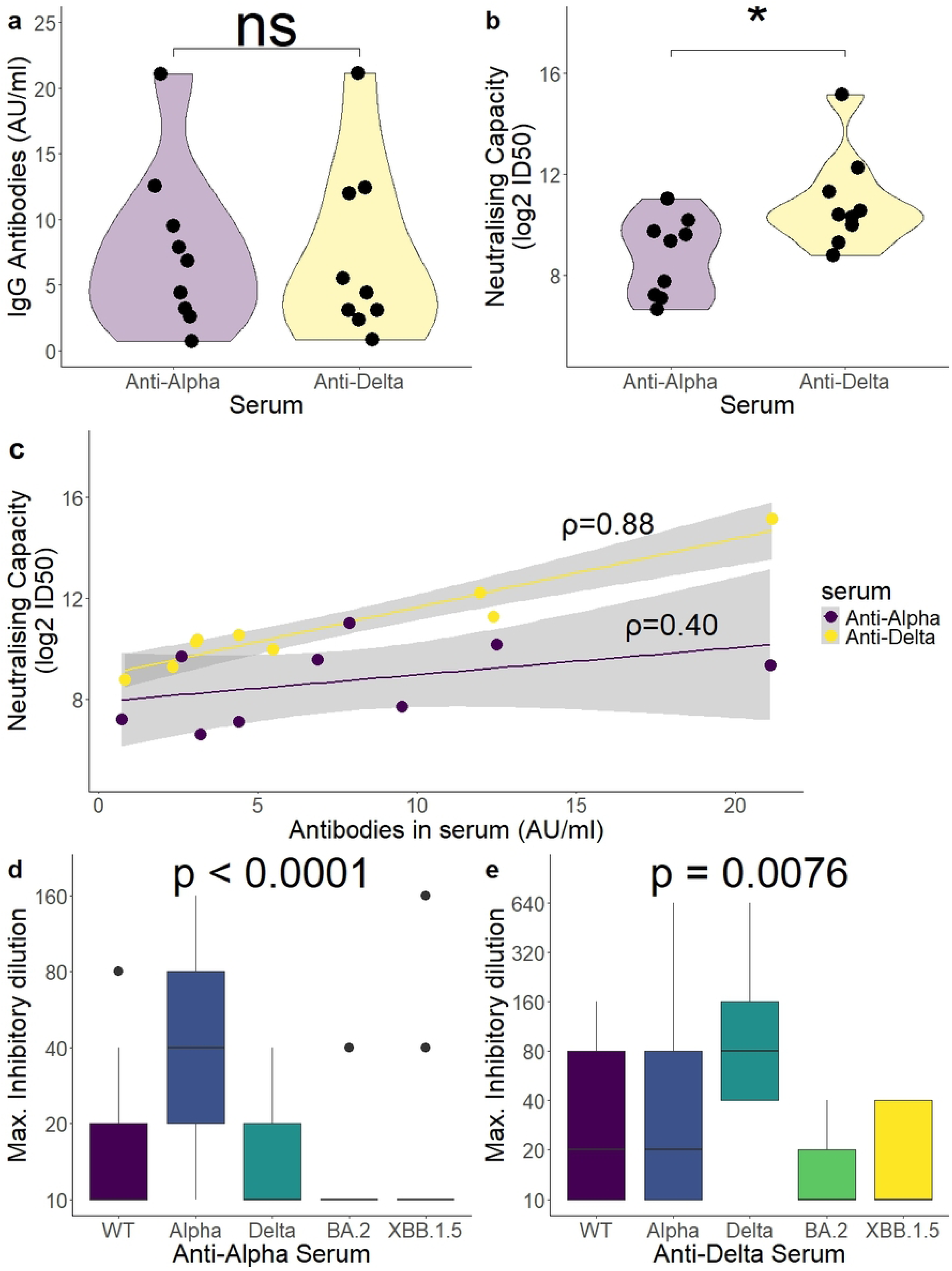
Neutralisation capacity post-infection is highest against variants of infection. **a)** 18 sera samples were selected for pseudovirus neutralisation, 9 from patients infected with Alpha variant and 9 infected with Delta variant, with paired IgG CLIA results (*p* = 0.73). **b)** Neutralising capacity of Delta-spike carrying pseudovirus particles. Anti-Delta sera were more successful at neutralising Delta pseudovirus than Anti-Alpha (*p* = 0.024) **c)** The CLIA assay is better predicting Delta neutralisation capacity in Anti-Delta than in Anti-Alpha sera (p = 0.003 *vs.* p = 0.29). **d,e)** Live virus neutralisation assay shows best neutralisation capacity for variant of infection, and a significant drop in neutralisation of Omicron variants. Minimum sera dilution tested is 1:20, with a value of 10 being indicated for samples with no neutralisation detected. For a) and b), Wilcoxon signed-rank test. For c), Spearman’s rank correlation coefficient. For d) and e), Kruskal-Wallis test. All data used to generate this figure is available in S4 File.

Despite paired antibody values measured *in vitro* (Fig 3A, *p* = 0.73), antibodies generated by individuals infected with the Alpha variant (anti-Alpha) performed significantly worse against the Delta variant lentivirus than the antibodies generated against the Delta variant (anti-Delta) (log2 ID50: 8.72 ± 1.57 (alpha) *vs.* 10.88 ± 1.89 (delta); *p* = 0.024) (Fig 3B). Notably, prediction of Delta neutralisation capacity by the *in vitro* assay (CLIA) was better for anti-Delta antibodies than for anti-Alpha, with Anti-Alpha CLIA results showing no significant correlation with Delta neutralisation capacity (Anti-Alpha ρ = 0.40, *p* = 0.29; Anti-Delta ρ = 0.88, *p* = 0.003; Spearman’s rank correlation coefficient) (Fig 3C).

### Live virus neutralisation assay shows a drop in Omicron neutralisation capacity for sera generated against pre-Omicron Variants of Concern (VOCs)

Neutralisation capacity was also measured using a live virus assay. In addition to the 18 samples tested with the pseudovirus assay, additional samples were added from the available Alpha group, all from 1-month after infection. Sera samples from Anti-Alpha (n = 21) and Anti-Delta (n = 9) were tested against viral variants wild-type (WT), Alpha, Delta, BA.2 (Omicron) and XBB.1.5 (Omicron). The neutralisation capacity was measured as the maximum inhibitory sera dilution where cytopathic effect was still observed. Comparing all variants tested, both sera performed best against their own variant (Alpha and Delta, Fig 3D-E). Anti-Alpha sera performed significantly better against Alpha than against all other variants tested, while Anti-Delta sera performed significantly better against Delta than against both Omicron variants tested (Dunn’s test, S5 File).

## Discussion

Our study shows a strong and variant-specific neutralising immune response after mild SARS-CoV-2 infection. There was an increase of the IgG response after one dose of vaccine after an infection period as measured by CLIA. Past studies have shown that one dose of vaccine in previously infected individuals (hybrid immunity) gives a higher IgG response than two doses of vaccine in covid-naïve individuals, suggesting the role of infection for priming the immune system [27–29].

Duration of the IgG response after vaccination is still a relevant issue during the SARS-CoV-2 vaccination campaigns, as the half-life of antibodies in blood is important for vaccination booster scheduling. One study showed a 50% decline of antibodies in a 1-3 month period after second dose, even in the case of pre-exposure to infection [30]. Similarly, in this study, post-infected patients with one dose of vaccine showed a mean decrease of 53% of igG levels three months after immunisation (22,4 ± 12,07 in month 3; 10,4 ± 6,71 in month 6 (mean ± IQR)). Notably, the IgG levels in month 6 showed comparable levels to the IgG levels prior to vaccination (p = 0.187). This steep decrease did not happen in the post-infection antibodies.

IgG levels are not the only relevant measure in humoral immunity, as different antibody fractions have different neutralisation capacities against different variants. It has been suggested that up to 66% of antibodies from a convalescent response are not cross-reactive, and are only able to neutralise one variant [40]. While the immunoassays will only give one readout as output, the neutralising antibody assay can be used to challenge sera against different viral variants. It is key to understand if immunoassays are still accurate even when measuring against emerging clades, especially as some immunoassays have been associated with reduced sensitivity to antibodies generated after Omicron infection, due to the antigens used in these kits being not updated to current circulating variants [31,32].

Accurately quantifying immune boosting has been specially relevant in the last vaccination campaigns with bivalent vaccines Wuhan/BA.5 (2022 campaign) and monovalent XBB.1.5 (2023 campaign). This is because some publications have suggested that, after booster vaccination with Omicron, the memory B cell response is directed towards epitopes of previous variants, decreasing the effect of boosting against new circulating variants (immune imprinting) [33–35].

A majority of individuals in the population have had their first exposure to SARS-CoV-2 in the Wuhan D614G variant, either due to the first available vaccines targeted against this variant or due to infection in the period before the variants of concern arose. Because of this, many publications have shown that neutralising capacity is highest against the ancestral variants than against other exposure variants, even after bivalent vaccination [36–38], omicron breakthrough infection [36,37] or monovalent XBB.1.5 vaccination [39]. In all cases, the variant of first exposure had the highest neutralising capacity. Our data suggests that this is not only true in the ancestral variant, but also in Alpha or Delta. Vaccine-naïve individuals infected with the Alpha variant showed the highest neutralisation against the same variant, and the same happened with individuals first exposed to the Delta variant (Fig 3D-E).

Regarding the Omicron variant landscape, neutralisation was significantly lower against Omicron variants than against the variant of exposure, including in XBB.1.5, the vaccine variant of the last season (Fig 3C-D). It is challenging to compare neutralisation against BA.2 and XBB.1.5 in our samples, as in both cases most sera samples fell below the limit of detection of neutralisation. Regardless, our data shows that antibodies targeting Alpha and Delta epitopes, perhaps more uncommon in the population as there have not been vaccination campaigns targeting these variants, show little neutralisation capacity against Omicron variants, at least after one exposure. A recent study studying the polyclonal responses from convalescent individuals found no nAbs with cross-reactivity between Delta and Omicron, which could explain the poor performance of Anti-Delta sera against the Omicron variants tested in our study [40]. Neutralising Abs for one SARS-CoV-2 variant can even be enhancing for other variants, at least *in vitro* [41].

Our data relating CLIA results to neutralisation data point that immunoassay values must be taken with caution when used in the clinic, as the ability to predict neutralising capacity from CLIA results depends on variant of first and second exposure. Remarkably, CLIA results and neutralisation capacity of the Delta variant did not significantly correlate in anti-Alpha sera samples, while showing a good correlation in anti-Delta sera samples (Fig 3C). While this publication has a small sample size in neutralisation assays, our results still show that precaution must be used when using IgG *in vitro* values to make clinical decisions.

During the COVID-19 pandemic, CLIA results have been used in the clinical context, such as in the detection of hyperimmune sera for its use as passive immunity [42]. If CLIA results are to be accurately used in the clinic, there is a need to obtain high-throughput *in vitro* assays with reagents that can be variant-specific, in order to better estimate immunity to a specific variant. This is even more relevant in the current context of SARS-CoV-2, where most individuals in the population have been exposed to various variants via both natural infection and vaccination.

Our data suggests a long period of viral shedding where RNA is still detectable, with the mean week of the first full negative sample being 5.20 ± 2.37. This is in contrast to other publications, where the median of viral RNA shedding to be around 17 days [1]. This is likely due to methodological differences. In the case of our study, no patient was considered negative until all three targets were negative, with a maximum CT value of 40. Other studies on viral shedding have used various targets and tests and used CT 40 as threshold [43], CT 38 as threshold [7] or CT 33 as threshold [44] amongst others, with varied results. Notably, our study measured one sample per week, as per the normal testing period in mild individuals during their isolation period in Spain. Daily testing, as performed in multiple studies with hospitalised patients, allows faster detection of negativity due to a higher number of datapoints. Isolation periods and patient release can be optimised with increasing testing, but this can be challenging for healthcare services working with mild patients isolated in their own place of residence.

The duration of infectious virus shedding has also been found to be affected by variables such as age, gender or viral variant. In our study, older participants had a higher viral load, even being individuals with mild disease (*p* = 0.008). There was no difference in shedding duration (*p* = 0.120). However, our study focused on adults between 18-55 year-olds. Elderly participants were discarded from inclusion as they were entering the vaccination campaign during the time of recruitment. A different study found faster viral clearance in younger participants, but only in individuals under 18 [7]. Additionally, males have been shown slower shedding than females in severe cases [8]. Our data, in mild patients, shows no difference in viral shedding by sex (*p* = 0.194). These variations highlight the complexities to determine, without using culture assays, when a patient can be released from a confinement without the risk of expansion of the virus.

In order to quantify samples in a more effective manner, our samples were also measured by dPCR. dPCR can more accurately quantify viral RNA than qPCR, as it can provide absolute quantification by partitioning the RT-PCR reaction mixture in nanowells containing, ideally, 0 or 1 target molecules, improving accuracy compared to qPCR [45,46]. From a clinical perspective, it provides the advantage of working without the need of RNA extraction, as partitioning the sample would also partition inhibitors present in the sample [47,48]. Avoiding the RNA extraction step could allow fast diagnosis in cases of emergency. Our data also supports the use of direct quantification for dPCR, as we found high correlation in the results between direct quantification and dPCR on an RNA extract (S1 Fig).

The correlation between dPCR and RT-PCR was high (R^2^ = 0.87), except for samples of low viral load, below the estimated limit of quantification for RT-PCR and dPCR [49,50]. Although dPCR can improve detection when the target concentration is low (above 30 ct) [11], both techniques show high quantification uncertainty in such cases, mainly due to undersampling errors [23,51].

Our data shows that, in mild covid, after week one, viral load is mostly residual, and probably associated with shedding of non-infectious viral RNA [52]. Therefore, late-CT positive results could be irrelevant in the context of isolation measurements, as CT value correlates inversely with probability of infectiousness [53]. We measured sgRNA in an additional RT-PCR assay, as it has been proposed as a good biomarker of active infection [5,26,49,54,55]. Nevertheless, it must be noted that another publication found sgRNA to have a poor yield as a measure of infective virus [56]. It has also been suggested that the decay ratio of sgRNAs after inhibition of transcription is not fast enough for sgRNAs to be an accurate predictor of infectivity [57].

In our cohort, a majority of individuals showed their first negative sgRNA result between week 1 and week 2 (Fig 1E), which is a timeframe similar to that expected for infectivity [56,58]. This shows the likelihood of sgRNA being a more accurate measurement of active infection than genomic RNA, and therefore it could be implemented as a diagnostic tool to avoid increasing isolation periods or hospital bed occupation. Notably, some studies have shown that while presence of sgRNA may not indicate infectiousness, probably due to the stability of the molecules in clinical samples, the absence of sgRNA would indicate absence of infectivity. Therefore, sgRNA has a high sensitivity as a marker of infectiousness despite its specificity being suboptimal [26,56]. In this model, a negative RT-PCR could be enough to consider an individual non-infectious.

In summary, this study sheds some light on SARS-CoV-2 shedding and immune response in groups that have been understudied, mild patients with pre-omicron VOCs (mainly Alpha and Delta variants). A majority of the studies of infection dynamics have been focused in hospitalised patients due to sampling being more accessible to healthcare professionals [1]. Additionally, most of the population was first exposed to ancestral variants either due to infection or vaccination.

The study has limitations: infectivity of clinical samples was not measured in culture, and the neutralisation sample number is low due to the low variability of variant circulation at the time of recruitment (69.5% patients infected with Alpha variant). Nonetheless, the study shows the limitations of accurately quantifying samples of late CT value, as well as a long period of detectable genomic RNA but absence of sgRNA, suggesting periods of no infectivity that should guide future strategies in releasing individuals from isolation or even to receive immunosuppressive therapies. Additionally, the data shows the low accuracy of commercial immunoassays in prediction of the neutralising capacity when the variant of first exposure and the variant of second exposure do not match, suggesting the need of updating reagents. These conclusions can guide diagnostic, pandemic response and vaccination schemes in future times of healthcare crisis.

## Materials and methods

### Patient selection and sample collection

A total of 105 patients diagnosed with infection by SARS-CoV-2 were recruited for the study. Ten participants dropped out of the study during the follow-up (Fig 4).

**Fig 4.**
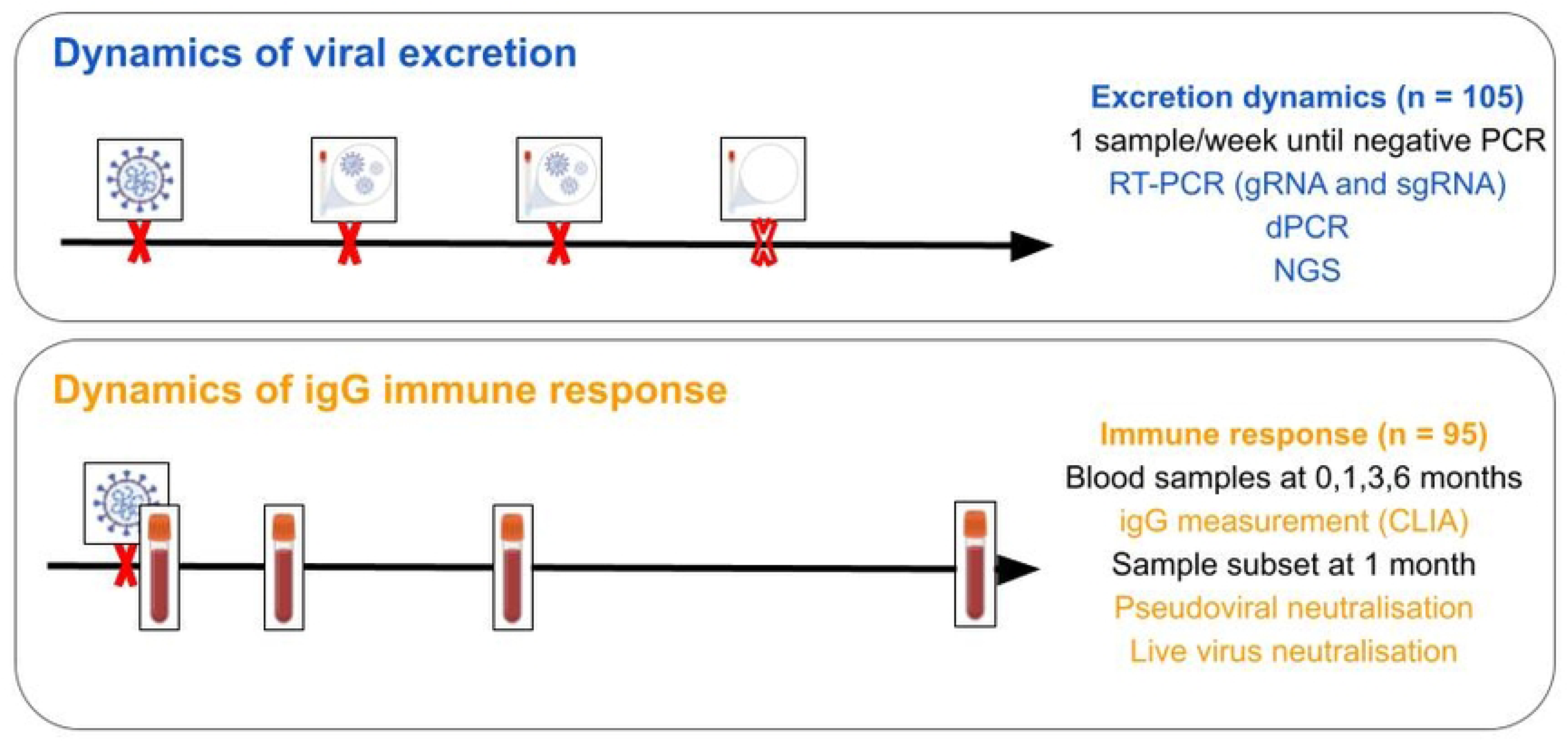
Graphical summary. For the first aim of the study, viral load was determined in one sample per participant and week until negative on three targets of an RT-PCR. For the second, serum samples were taken at 0, 1, 3 and 6 months after infection in order to measure IgG levels and study neutralisation capacity.

For the measurement of viral dynamics, one nasopharyngeal sample was taken (Viral transport medium, Vircell, Granada, Spain) per week until the virus was undetectable (negative RT-PCR for all three targets). For the study of the immune response dynamics, a serum sample was taken from patients at months 0, 1, 3 and 6 after infection in order to measure antibody levels. As participants were non-hospitalised and quarantined in their places of residence, a healthcare professional was in charge of visiting participants and obtaining the samples until they could go to the Hospital for the sample collection.

The study received approval from the Galician Network of Research Ethics Committees (protocol number 2020/627), adhering to the principles outlined in the Declaration of Helsinki. All methodologies were conducted in accordance with relevant guidelines and regulations, and all participants provided informed consent prior to their inclusion in the study.

### Inclusion criteria

Non-hospitalised patients aged 18-55 years, with laboratory-confirmed SARS-CoV-2 infection by RT-PCR in the recruitment period (March - July 2021). All participants signed an informed consent form.

### Viral RNA quantification

#### RT-PCR

Nucleic acid extraction was performed on a Microlab STARlet IVD platform (HAMILTON) using the STARMag 96×4 Universal Cartridge Kit (Seegene Inc., Seoul, Korea).. RT-PCR was performed with Allplex 2019-nCoV Assay, (Seegene Inc.) and the Bio-Rad CFX96 Touch™ (Bio-Rad Laboratories, Inc. USA), targeting the Envelope (E), Nucleocapsid (N) and RNA-Dependent RNA Polymerase (RDRP). An additional RT-PCR was performed to target the E gene sgRNA, using the primers and probe previously published by Wölfel *et al*. [5] and the Lightcycler Multiplex RNA Virus Master (Roche Diagnostics, Mannheim, Germany) on z480 thermocycler (Roche Diagnostics), with a final concentration of primers of 0.1 μM and probe of 0.05 μM. .

#### Digital PCR (dPCR)

Nasopharyngeal exudate crude samples were quantified (direct quantification) using the QiaAcuity One-step Viral RT-PCR kit from Qiagen (Qiagen, Hilden, Germany) with a 26k nanoplate following manufacturer’s specifications. The primer/probe used was the COVID-19 E-gene primer-probe reagent (TIBMOLBIOL, Berlin, Germany) with 1 μl per 40 μl reaction. Following recommendations from dMIQE (Minimum Information for Publication of Digital PCR Experiments) [23], only samples with over 15000 valid partitions and below 0,9 positive partition ratio were considered as valid. In order to avoid nanoplate saturation, samples were diluted based on the CT value obtained on the RT-PCR as follows: CT < 12 dilution 1:1000, CT = 12-14 dilution 1:200, CT = 14-15.5 dilution 1:100, CT = 15.5-16 dilution 1:50, CT = 16.5-18 dilution 1:20, CT 18-19 dilution 1:10.

### SARS-CoV-2 Next-Generation Sequencing

In order to know the viral variant, one sample per patient was selected for viral whole genome sequencing by NGS. For NGS, 11 μl of extracted RNA were subjected to retrotranscription using SuperScript IV kit (SSIV, Invitrogen by Life Technologies, Carlsbad, USA) according to manufacturer’s specifications and using random hexamers as primers. cDNA was then amplified using the Q5 High Fidelity DNA Polymerase (New England Biolabs, Ipswich, USA) and the ARTIC v3 primer panel. Enriched samples were then normalised and libraries were prepared using the Illumina DNA prep kit (Illumina Inc., San Diego, USA) using 1/4 of the recommended volume. The Illumina iSeq platform was used for sequencing. Fastqs generated were analysed as described in https://github.com/OMIC-G/COV and submitted to GISAID.

### Measurement of antibodies in sera

Sera samples were taken at months 0, 1, 3 and 6 after diagnosis and analysed using the COVID-19 VIRCLIA IgG Monotest (Vircell Microbiologists, Granada, Spain), a CLIA *in vitro* assay targeting IgG antibodies against the receptor binding-domain of the spike protein.

### Pseudovirus neutralisation assay

The protocol for the pseudovirus neutralisation assay was based on the previously published by Nie *et al.* [24]. Commercially available lentiviral vectors transfected with green fluorescent protein (GFP) gene and presenting SARS-CoV-2 spike protein variant Delta (B.1.617.2) were used (VectorBuilder, Neu-Isenburg, Germany). Sera was heat-inactivated (30 min, 56°C) and serially diluted (2X) in duplicates five times. The lentiviral vectors were added to the sera in a virus/cell ratio (Multiplicity of Infection, MOI) of 50. A positive control (no serum, viral control) and a blank (no virus, cellular control) were added on each plate in triplicate.

Incubation in 96 tissue culture-treated well plates was performed for 1 hour at 37°C. Then, HEK293T-ACE2 cells (25000 cells/well) were added and incubated for 72 hours at 37°C, 5% CO2 in a FLUOstar Omega Fluorescence plate reader (BMG LABTECH, Offenburg, Germany). After 72 hours, fluorescence was measured as the sera dilution that could reduce fluorescence to 50% of the viral control (no sera) (50% inhibitory dilution (ID50)). ID50 was calculated using the Reed-Muench method.

### Live virus neutralisation assay

For the live virus neutralisation assay, heat-inactivated sera (30 min, 56°C) was serially diluted (2X) eight times starting at 1:20 dilution, and then incubated for 1 hour at 37°C with viral samples diluted to a viral infectivity of 100 times the median tissue culture infective dose (100x TCID50).

The viral variants used for neutralisation were wild-type (Ancestral), Alpha (B.1.1.7), Delta (B.1.617.2), Omicron (BA.2) and Omicron (XBB.1.5). After incubation, the mix was added to a 96-well plate seeded with 25000 Vero cells/well. After five days of infection, maximum inhibitory concentration was calculated as the maximum sera dilution with no observed cytopathic effect. An infection control (no sera) and a positive control (high-titer sera) were added in each experiment.

## Data analysis

All statistical data analysis was done using R (version 4.1.1, https://cran.r-project.org/). Data visualisation was performed with the R program ggplot2 [25]. Shapiro-Wilk normality test was performed to check for normality. Kruskal-Wallis and Dunn’s tests were used when indicated. When multiple pairwise tests were performed simultaneously, *p*-value was adjusted using the Holm correction. Pearson’s correlation coefficient (R^2^) and Spearman’s rank correlation coefficient (ρ) was used for rank correlations between two variables when indicated.

## Acknowledgements

The live virus neutralisation experiments were performed in the BSL-3 laboratory in the installations of Vircell S.L (Granada, Spain). We would like to thank their support and expertise in viral culture and neutralisation.

We would like to thank all participants for volunteering to the development of this study.

We would like to acknowledge all workers in the Complexo Hospitalario Universitario de Vigo (CHUVI) for their dedication and passion in their work, and Fátima Mella Castro for her work obtaining the biological samples needed for this study.

## Supporting Information

**S1 Fig.**
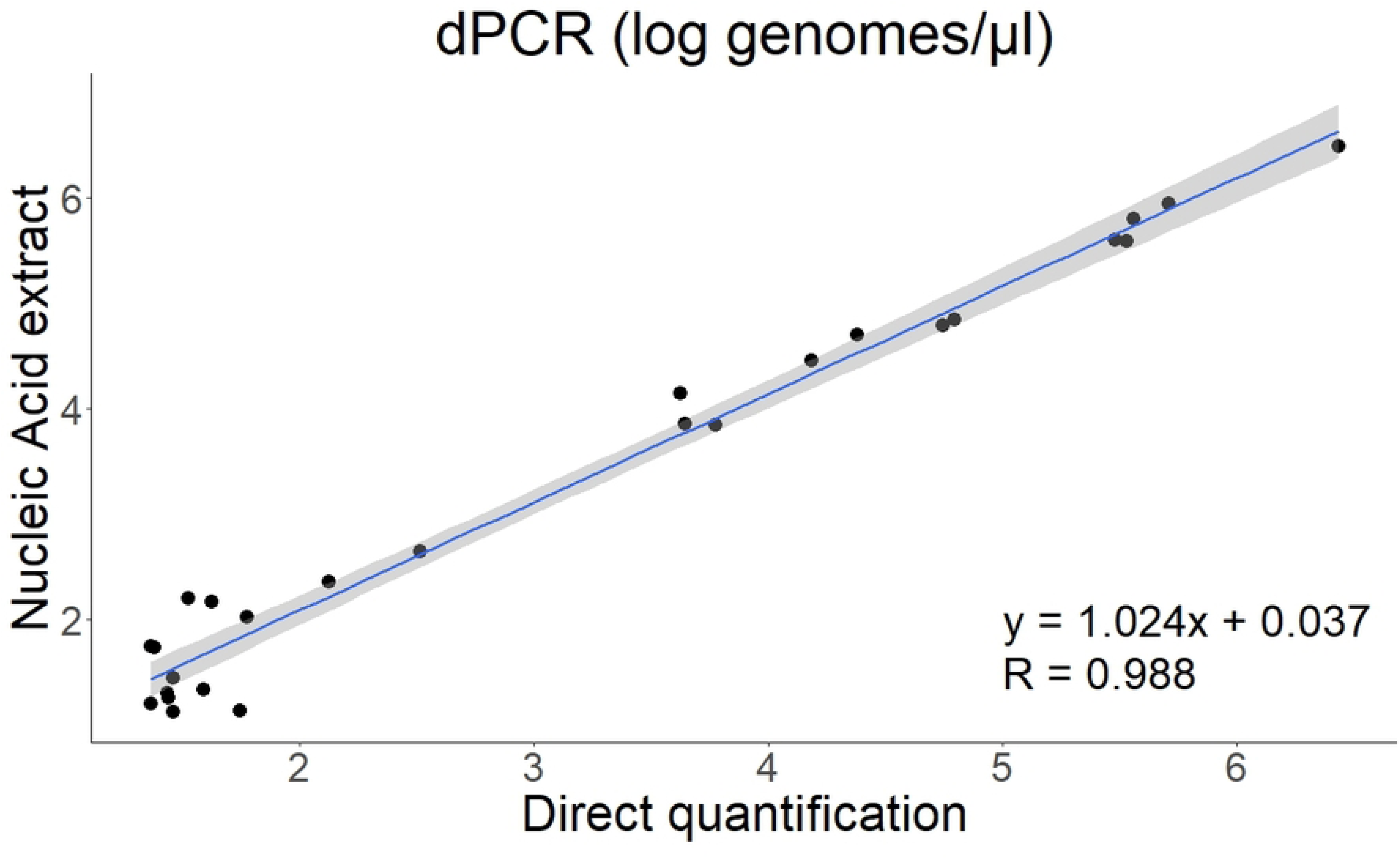
dPCR can be effectively used for viral load quantification without an extraction step. In order to ensure the reliability of the direct quantification compared with the RNA extract, a subset of 27 samples was quantified using both an RNA extract and the crude sample. The comparison shows no difference in the results of the digital PCR (R^2^ = 0.988), suggesting that nucleic acid extraction is an unnecessary step in SARS-CoV-2 quantification via dPCR.

**S2 Fig.**
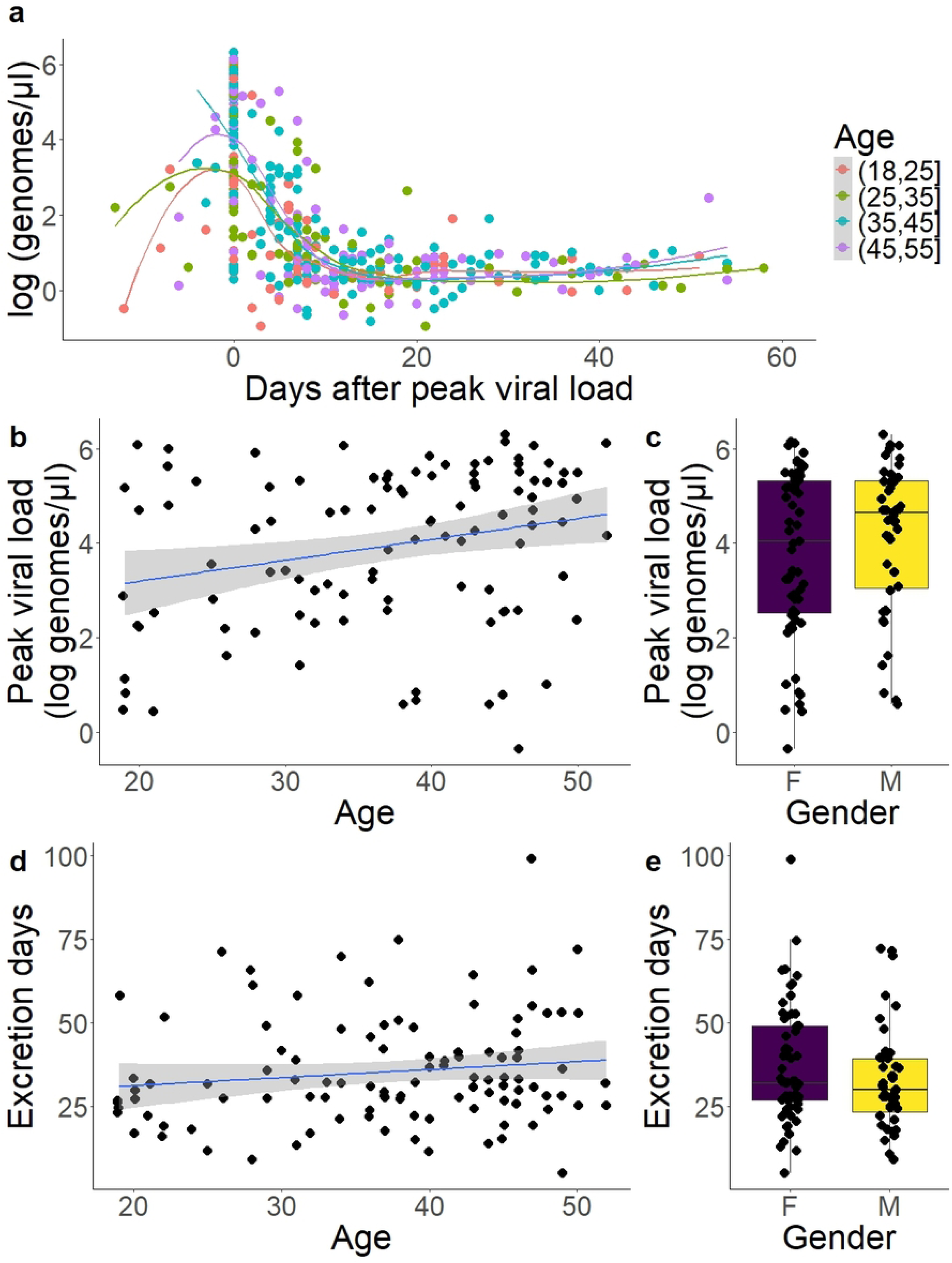
Impact of age and gender in excretion dynamics. **a)** Excretion dynamics plot colored by age. Peak viral load (x = 0) is higher in older groups. **b)** There is a mild correlation between age and peak viral load (*p* = 0.008) **c)** Gender did not significantly impact peak viral load (*p* = 0.367). **d)** There was no statistically significant correlation between age and the number of excretion days (days with viral RNA detectable by RT-PCR) (p = 0.120) **e)** Gender did not significantly impact the number of excretion days (p = 0.194). For b) and d), Spearman’s rank correlation coefficient. The gray area represents 0.95 confidence interval. For c) and e), Wilcoxon Signed-rank test.

**S1 File. Raw data from shedding dynamics analysis (Fig. 1)**

**S2 File. Statistical analysis results from igG measurement data (Fig 2).**

**S3 File. Raw data from igG measurement data (Fig. 2)**

**S4 File. Raw data from neutralisation capacity analysis (Fig. 3)**

**S5 File. Statistical analysis results from live virus neutralisation (Fig 3D,E).**

